# A seasonal pulse of ungulate neonates influences space use by carnivores in a multi-predator, multi-prey system

**DOI:** 10.1101/2021.12.16.472870

**Authors:** Joel Ruprecht, Tavis D. Forrester, Nathan J. Jackson, Darren A. Clark, Michael J. Wisdom, Mary M. Rowland, Joshua B. Smith, Kelley M. Stewart, Taal Levi

## Abstract

1. The behavioral mechanisms by which predators encounter prey are poorly resolved. In particular, the extent to which predators engage in active search for prey versus incidentally encountering them is unknown. The distinction between search and incidental encounter influences prey population dynamics with active search exerting a stabilizing force on prey populations by alleviating predation pressure on low-density prey and increasing it for high-density prey.
2. Parturition of many large herbivores occurs during a short and predictable temporal window in which young are highly vulnerable to predation. Our study aims to determine how a suite of carnivores responds to the seasonal pulse of newborn ungulates using contemporaneous GPS locations of four species of predators and two species of prey.
3. We used step-selection functions to assess whether coyotes, cougars, black bears, and bobcats actively searched for parturient females in a low-density population of mule deer and a high-density population of elk. We then assessed whether searching carnivores shifted their habitat use toward areas exhibiting a high probability of encountering neonates.
4. None of the four carnivore species encountered parturient mule deer more often than expected by chance suggesting that predation of young resulted from incidental encounters. By contrast, we determined that cougar and male bear movements positioned them in proximity of parturient elk more often than expected by chance which is evidence of searching behavior. Although both male bears and cougars searched for neonates, only male bears used elk parturition habitat in a way that dynamically tracked the phenology of the elk birth pulse suggesting that maximizing encounters with juvenile elk was a motivation when selecting resources.
5. Our results support the existence of a stabilizing mechanism to prey populations through active search behavior by predators because carnivores in our study searched for the high-density prey species (elk) but ignored the low-density species (mule deer). We conclude that prey density must be high enough to warrant active search, and that there is high interspecific and intersexual variability in foraging strategies among large mammalian predators and their prey.

## INTRODUCTION

Although the distinction between active search and incidental encounter of prey is key to understanding predator-prey dynamics, whether, and under what conditions predators actively search for particular prey species is poorly understood. Diet breadth theory suggests that predators seeking to maximize their rate of energy acquisition should have a more generalist prey profile until the encounter rate with the most profitable prey item crosses a critical threshold that makes it suboptimal to include lower-profitability prey in their diet (MacArthur & Pianka 1966; Figure 1; Supporting Information Text S1). Diet specialization on abundant prey is necessarily associated with active search for that prey, but as the density of the focal prey species declines, additional species enter the diet which reduces the rate of acquisition of the focal prey. A switch from active search for a prey species at high density to incidental encounter of the same species at low density can have stabilizing effects on prey population dynamics (Murdoch 1969) and results in the canonical sigmoid Type III functional response that is associated with many of the most interesting dynamics in predator-prey systems including alternative stable states, bifurcations, and predator pits (Holling 1959; Messier 1994; Levi *et al.* 2015).

**Figure 1:**
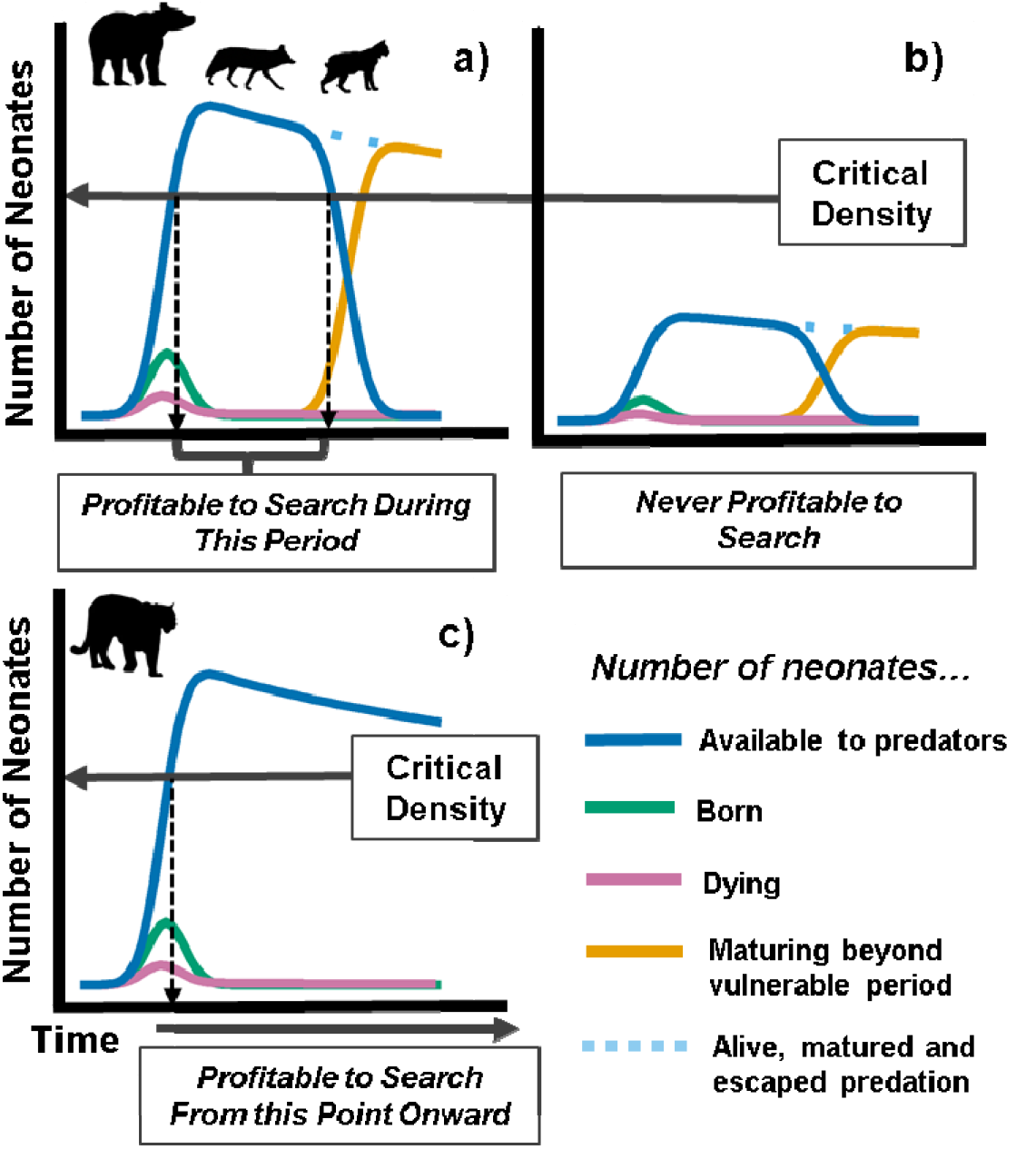
Conceptual diagram showing birth pulse phenology and the profitability of predator search behavior. Panels a) and b) are representative of species that can only successfully capture neonates during a limited time period (e.g., bears, coyotes, bobcats), beyond which neonates mature to the point where they have escaped the majority of predation. Panel a) represents a population with a high density of neonates; the period in which the number of neonates available (blue line) exceeds the critical density is the period when active search is profitable. Panel b) represents a population with a low density of neonates that never exceeds the critical density such that actively searching is never the optimal strategy. Panel c) represents species that can capture juvenile ungulates throughout their entire first year of life (and beyond; e.g., cougars), such that it is profitable to search as soon as the critical density has been exceeded.

The density-dependent response of predators that is evidence for active search can be found by observing disproportionate predation in years in which a given prey species is at high density. However, in many systems prey densities vary predictably within a year due to a birth pulse that provides a uniquely vulnerable life stage for predators. A classic case of this phenomenon is the seasonal pulse of songbird nests (Schmidt, Goheen & Naumann 2001; Vigallon & Marzluff 2005), and both active search (Pelech, Smith & Boutin 2010) and incidental encounter (Schmidt 2004) strategies have been documented as a function of nest density, vulnerability, and maternal vigilance or defense (Schmidt 1999). In addition to songbirds, large herbivore systems worldwide feature a predictable birth pulse of neonates that are vulnerable to predators across a wide body size gradient. Although mortality of ungulate neonates has been studied extensively (Linnell, Aanes & Andersen 1995), it is still largely unknown whether, and under what conditions predators engage in active search behavior during the birth pulse, which has important implications for prey population dynamics.

The propensity of carnivores to target neonates likely depends on factors intrinsic to both the predator and prey. For predators, this may vary by body size, hunting proficiency or mode, and ability to cope with maternal defense by the large herbivore. For example, ungulate neonates are highly vulnerable to predation immediately following parturition from a suite of carnivores, but quickly become sufficiently vagile to elude those that are less predaceous or of smaller body size. Thus, species such as bears and many mesopredators experience a particularly short resource pulse of neonates (Linnell, Aanes & Andersen 1995; Zager & Beecham 2006; Griffin *et al.* 2011) (Figure 1a, b). In contrast, the birth pulse may be less consequential to the largest felids and canids that can efficiently capture large-bodied ungulates throughout the first year of life and beyond (Figure 1c). Even within a species, the response of predators to the ungulate birth pulse may vary by individual or sex as has been shown in bears (Jacoby *et al.* 1999; Zager & Beecham 2006; Rayl *et al.* 2015).

Because predatory behavior during the ungulate birth pulse is influenced by idiosyncratic combinations of factors intrinsic to both predators and prey, studies involving multiple carnivore and multiple ungulate species will be needed to elucidate the generality of search behavior. Previous research has largely focused on identifying spatial shifts in habitat use by bears (but see Bastille-Rousseau *et al*. 2016; Svoboda *et al*. 2019) toward birthing grounds or areas on the landscape more likely to contain neonates, which has resulted in different conclusions wherein both active search (Rayl *et al*. 2018) and incidental encounter (Bastille-Rousseau *et al.* 2011; Bowersock *et al.* 2021) were inferred. Much stronger inference is possible using analytical methods that utilize spatiotemporal encounters between individual predators and individual prey from contemporaneous GPS telemetry data. Further, identifying general conditions associated with incidental encounter and active search requires relocation data on multiple species of predators and prey across gradients in body size, abundance, and life histories.

Here we use contemporaneous GPS tracking data at the level of encounters between individual predators and individual prey to assess whether carnivores actively searched for or incidentally encountered ungulate neonates in a multi-predator, multi-prey system. Each carnivore species (cougar [*Puma concolor*], coyote [*Canis latrans*], black bear [*Ursus americanus*], bobcat [*Lynx rufus*]) varied in size, life history, and predatory ability, ranging from large-bodied obligate predators to smaller-bodied omnivores, while prey species (mule deer [*Odocoileus hemionus*] and elk [*Cervus canadensis*]) varied dramatically in abundance. Our primary objective was to determine whether predators encountered parturient female ungulates more often than expected by chance, indicating active search, while controlling for their habitat preferences. If predators did indeed exhibit targeted search behavior, a secondary objective was to determine whether a shift in space use toward parturition habitat tracked the phenology of the birth pulse consistent with an effort to maximize detections of neonates. We hypothesized that the magnitude of response by predators to the birth pulse would be greater toward elk than mule deer for all carnivore species because elk were approximately 5 times more abundant than mule deer and could thus cross the profitability threshold associated with specialization under diet-breadth theory. We expected the carnivore species with a more generalist diet profile (bears and coyotes) would alter their foraging behaviors more strongly than cougars or bobcats because generalist consumers are more fluid in their response to changing resources (Ostfeld & Keesing 2000; Yang et al. 2008). Bears are the least carnivorous of these taxa, and previous research suggests that male bears are disproportionately predaceous (Rode, Robbins & Shipley 2001, Boertje *et al.* 1988; Jacoby *et al.* 1999), so we additionally hypothesized that male bears would be more likely to exhibit active search behavior. Further, cougars kill mule deer and elk of all age classes year-round (Clark *et al.* 2014), so we expected that the ungulate birth pulse may be less consequential to cougars given their adeptness at killing larger prey such that they may not exhibit a change in behavior during the earliest neonatal period. Finally, we hypothesized that bobcats would show the weakest response, since they rarely consume elk and mule deer in our study area (Ruprecht *et al.* 2021b).

## MATERIALS AND METHODS

### Study area

Our study was conducted in the Blue Mountains of northeastern Oregon centered at the Starkey Experimental Forest and Range between 2016 and 2019 (Figure 2). The major habitat types in the study area include grasslands, riparian areas, open forest dominated by ponderosa pine (*Pinus ponderosa*), and closed forest consisting of a mixture of Douglas fir (*Pseudotsuga menziesii*), grand fir (*Abies grandis*), larch (*Larix occidentalis*), ponderosa pine and lodgepole pine (*Pinus contorta*). Riparian areas sustain willows (*Salix* spp.), hawthorn (*Crataegus* spp.), Rocky Mountain maple (*Acer glabrum*), and other shrub species in low quantities. Starkey Experimental Forest and Range and adjacent public lands support an assemblage of native and domestic large herbivores including mule deer, white-tailed deer (*Odocoileus virginianus*), elk, and seasonally grazed domestic cattle (*Bos taurus;* Rowland *et al.* 1997). Carnivore species include black bears, coyotes, cougars, and bobcats. Gray wolves (*Canis lupus*) are colonizing the area but currently occur only occasionally and unpredictably in the study area.

**Figure 2:**
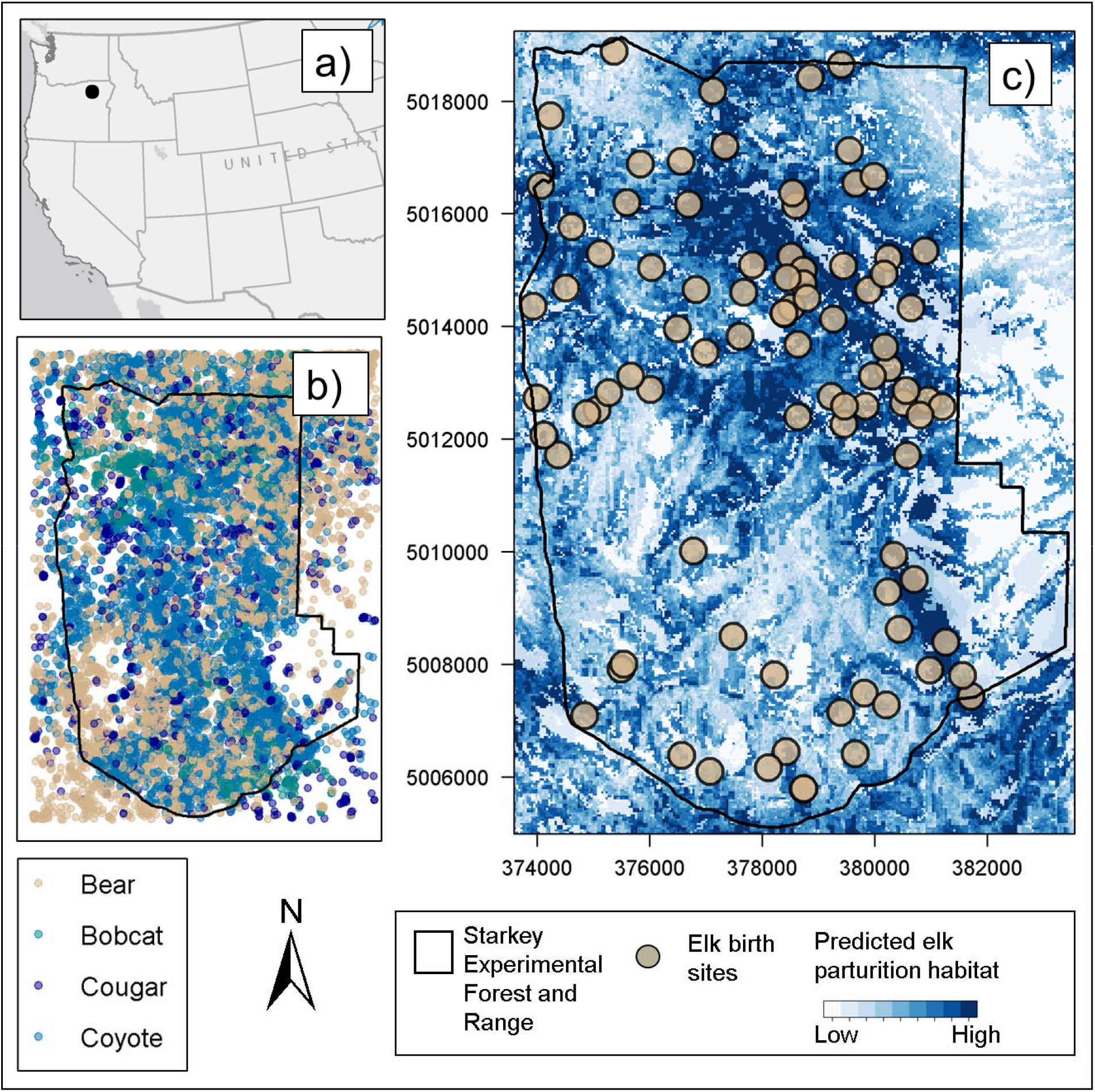
Study area: a) the general location of study area in northeast Oregon, USA, is indicated by the black dot. b) GPS telemetry locations of carnivores (black bears: tan points, bobcats: green points, cougars: dark blue points and coyotes: light blue points) used for analysis. c) Starkey Experimental Forest and Range boundary (black polygon) overlaid on the elk parturition habitat resource selection function (white areas indicate areas with low probability of selection, dark blue areas indicate high probability of selection). Tan points represent elk birth sites.

We used global positioning system (GPS) telemetry data from 9 cougars (representing 11 animal years), 17 coyotes (21 animal years), 11 black bears (18 animal years), 6 bobcats (7 animal years), 25 adult female mule deer (45 animal years), and 59 adult female elk (89 animal years). GPS positions were recorded every 2 or 3 hours for carnivores, every 30 minutes for elk, and every 60 or 90 minutes for deer. Details on capture and handling of carnivores, elk, and mule deer can be found in Ruprecht *et al.* (2021a), Wisdom *et al.* (1993) and Jackson *et al.* (2021), respectively. All animal capture and handling adhered to protocols approved by the USDA Forest Service, Starkey Experimental Forest Institutional Animal Care and Use Committee (IACUC No. 92-F-0004; protocol #STKY-16-01) and followed the guidelines of the American Society of Mammalogists for the use of wild mammals in research (Sikes 2016).

### Identifying elk and mule deer parturition events

We inferred elk parturition events from GPS-collared elk between 2016 and 2019 using the rolling minimum convex polygon (MCP) method described by Nicholson *et al.* (2019) to estimate large herbivore parturition events based on localized movements. We assigned a parturition event for elk as the first day a rolling MCP (based on a 24-hour window) decreased to <30 hectares or less for a minimum of 120 hours. If these conditions were not met, we assumed the elk did not give birth. We validated the method using independent elk GPS locations with known birth dates and locations (*N* = 30) from a previous study in the same area (Long et al. 2016). The mean discrepancy between birth dates determined from field investigations by Long et al. (2016) and birth dates predicted by the rolling MCP method was 18 hours.

Parturition events of mule deer were determined either by 1) monitoring GPS-collared adult females for localized movement and then searching the area for neonates where a cluster of GPS locations had formed, 2) using vaginal implant transmitters (details in Jackson *et al.* (2021)), or 3) using the rolling MCP inference method (Nicholson *et al.* 2019). For the rolling MCP method, we assigned a parturition event for deer as the first day a rolling MCP (based on a 24-hour window) was <15 hectares or less for a minimum of 120 hours. We validated the rolling MCP method for 14 deer parturition events determined from field investigations and found a mean discrepancy of 33 hours.

### Objective 1: determining whether predator movements indicated active search for parturient ungulates

We determined whether predators were actively searching for parturient female ungulates by assessing whether their actual movements (hereafter “steps,” or the Euclidean distance between subsequent GPS relocations) led to encounters with parturient females more often than hypothetical steps that they could have taken but did not. To do so, we fit step-selection functions (Fortin *et al.* 2005) for each carnivore species to estimate the relative probability of selecting a location on the landscape, given its previous location and a suite of covariates at the ending locations of both real and hypothetical (hereafter, “random”) steps. We excluded data from individuals whose home ranges did not overlap deer and elk relocations in Starkey Experimental Forest and Range. The covariate of primary interest was whether the endpoint of each observed or random step was within a 200-meter proximity of a parturient female ungulate. By comparing whether the observed steps were more often within close proximity of a parturient ungulate than were random steps, we could assess whether predators detected neonates more often than expected by incidental encounter, which is evidence of active search behavior. We restricted the analysis to include location data of each female ungulate in the 30 days after a parturition event was predicted to occur and fit separate models for deer and elk for each of the four carnivore species (8 models total). We compared each real step (coded 1) with 20 random steps (coded 0; Latombe *et al*. 2014) and fit models using conditional logistic regression. The random locations were generated by taking random draws from the fitted distributions of step lengths and turning angles (Gamma and von Mises distributions, respectively; Avgar *et al.* 2016) constructed from GPS data for each predator species and projecting the random locations for each GPS position onto the landscape given the previous location. To control for the possibility that observed steps landed in close proximity to an ungulate simply because of the predator’s preference for certain landscape or vegetative features, we included additional covariates known to influence carnivore movements (Ruprecht *et al.* 2021b): canopy cover (derived from LEMMA’s generalized nearest neighbor model; Ohmann *et al.* 2011), potential vegetation type (a factor variable with classes for open forest, closed forest, grassland, and other), ruggedness (using the vector ruggedness measure, a composite index of terrain encompassing both slope and aspect; Sappington, Longshore & Thompson 2007), the distance to nearest open road (natural log transformed), and distance to nearest perennial water source (natural log transformed). All continuous variables were centered to have a mean of 0 and scaled to have a standard deviation of 1. The step-selection function took the form *w(x)* ~ *exp*(*β*_1_ × parturient deer or elk presence + *β*_2_ × canopy cover + *β*_3_ × potential vegetation type + *β*_4_ × ruggedness + *β*_5_ × *ln*(distance to road) + *β*_6_ × *ln*(distance to perennial water source) + *β*_7_ × *ln*(step length) + *β*_8_ × *cosine*(turning angle)). Because the response to neonates can differ by sex of bears (Rayl *et al.* 2015), we fit additional models for male and female bears separately. We did not fit additional sex-specific models for cougars, coyotes, or bobcats, however, because we had no evidence *a priori* that predation of neonates would differ between the sexes for those species.

### Objective 2: determining whether use of parturition habitat by searching predators tracked the phenology of the birth pulse

One strategy for predators to maximize encounters with neonates is to shift habitat use to areas with a high probability of selection by parturient female ungulates. To this end, we constructed a resource selection function (RSF; Manly *et al.* (2007)) using GPS locations of adult female elk in the 7 days immediately following a parturition event with landscape and vegetative characteristics hypothesized to influence location of parturition sites in our area based on previous research (Johnson *et al.* 2000, Stewart *et al.* 2002, Long, Rachlow & Kie 2008). The resource selection function for elk parturition habitat took the form *w(x)* ~ exp(*β*_1_ × canopy cover + *β*_2_ × *ln*(distance to open road) + *β*_3_ × *ln*(distance to perennial water source) + *β*_4_ × ruggedness + *β*_5_ × shrub cover + *β*_6_ × forb cover + *β*_7_ × slope + *β*_8_ × aspect + *β*_9_ × elevation + *β*_10_ × potential vegetation type), where *w*(*x*) is the relative probability of selection for adult female elk during the first 7 days following parturition. Percent shrub and forb cover variables were derived from LEMMA’s generalized nearest neighbor model (Ohmann *et al.* 2011); slope, aspect, and elevation were drawn from a digital elevation model; and other covariates are as described above. We paired each used elk location (coded as 1) with 10 randomly generated locations representing locations available to but not used by elk (coded as 0) and used a generalized linear model with a binomial error distribution and logit link to model the relative probability of selection. Given that our objective was to create a spatially-explicit map of the study area predicting the areas with a high probability of selection by parturient ungulates and not make inference on the specific resources that they selected for, we did not conduct model selection and instead used the global model for predictions. Each 30 × 30 m pixel in the resulting predictive map projected onto the study area represented *w*(*x*), or the RSF score for parturition habitat. We then calculated the mean value of the RSF scores across all GPS locations for each carnivore species that exhibited active search behavior (determined from the previous analysis) on a weekly basis from 15 April to 31 to determine whether predators shifted habitat use toward places likely to be inhabited by ungulate neonates. We predicted that predator use of habitat used for parturition by elk would decline later in the season if predators were attempting to maximize encounters with neonatal elk by shifting habitat use. Weekly mean RSF scores represented each predator’s use of predicted parturition habitat with higher values indicating higher use of parturition habitat, where “use” is a measure of the investment in a set of resource units by an animal during a sampling period (Lele *et al.* 2013).

We used the weekly average elk parturition RSF score at carnivore GPS relocations as the response variable in a generalized linear mixed model and used Julian week (i.e. the number of weeks elapsing since January 1) as a predictor to assess how carnivore use of elk parturition habitat changed throughout the season. We included a random intercept for animal ID to control for differences in the mean elk parturition RSF score available within individual predator home ranges. We fit models with both linear and quadratic effects of Julian week as predictors. We hypothesized that if a quadratic effect of Julian week on use of parturition habitat tracked the phenology of the birth pulse, it would be indicative of an effort of predators to alter habitat use to maximize encounters with neonates immediately following parturition. Alternatively, lack of a quadratic relationship between predator use of parturition habitat and Julian week could indicate the predator does not spend time in areas where neonates are likely to be immediately following parturition, or that the predator may actively hunt older, more mobile neonates or other age classes of prey. We used likelihood ratio tests to compare whether the quadratic effect of Julian week was supported over a linear effect.

## RESULTS

We identified 45 parturition events for deer (14 determined from field investigations and 31 determined using the rolling MCP method) and 89 parturition sites for elk (all determined using the rolling MCP method) between 2016 and 2019 (Figure 2c). We estimated 27 May (range: 10 May to 25 June) as the mean parturition date for elk and 02 June (range: 21 May to 28 June) as the mean parturition date for deer across all 4 years in our study.

### Step-selection functions to infer active search behavior

Of the four carnivore species, none made movements such that they encountered parturient mule deer (within a 200-meter proximity) in the first 30 days post parturition more often than the random movements generated in the step-selection function (Figure 3a; Figure 4a; Tables S1–S2) suggesting they were not actively searching for mule deer neonates. We documented no GPS-collared cougars or bobcats within 200 meters of a GPS-collared mule deer at the time a simultaneous fix was taken during the 30 days following parturition so models could not be fit for these species. For elk, only two carnivore species exhibited movements that positioned them in proximity of elk post-parturition more often than did hypothetical movements, cougars (β_elk presence_ = 0.94, *P* = 0.001; Figure 3b; Table S3) and bears, although for bears the result was marginally significant (β_elk presence_ = 0.30, *P* = 0.059; Figure 3b; Table S3). Sex-specific analysis revealed that male bears were more likely to encounter elk than expected by chance (β_elk presence_ = 0.56, *P* = 0.004, *N* = 7 bears; Figure 4b; Table S4) while there was no evidence supporting this effect for female bears (β_elk presence_ = −0.45, *P* = 0.24, *N* = 4 bears; Figure 4b, Table S4). Projecting model predictions onto the landscape spatially and overlaying GPS locations of post-parturient female elk in the 30 days post birth further revealed similarities between areas used by neonatal elk and areas selected for by male bears and cougars as documented by the high degree of spatial overlap (Figure 5a,c). Female bears, however, generally did not select for areas that overlapped with elk neonates (Figure 5b), suggesting their space use decisions were driven more heavily by other resources that were unrelated to elk.

**Figure 3:**
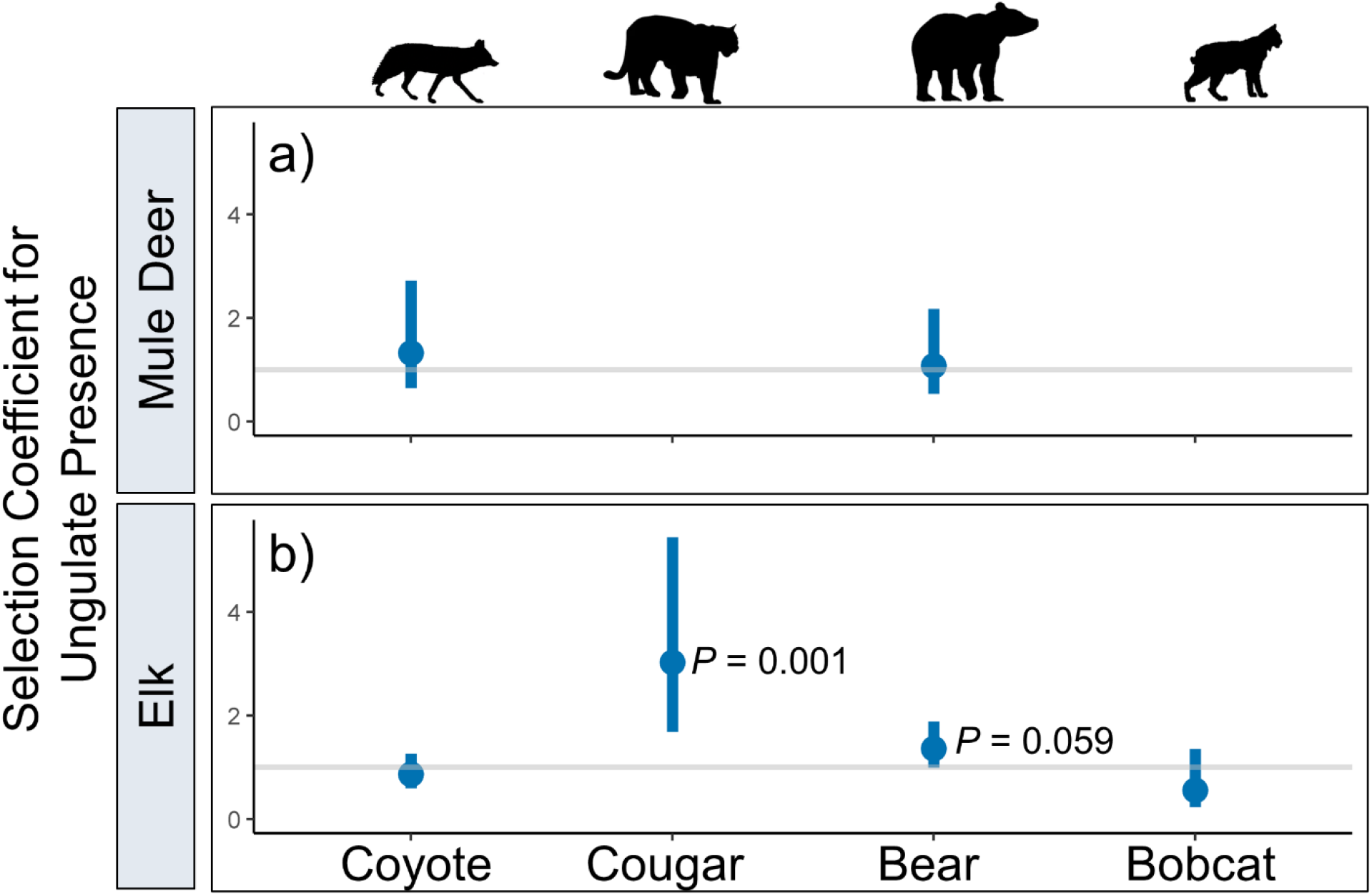
Selection coefficients (exp(β_ungulate presence_)) from step-selection functions assessing whether carnivores (coyotes, cougars, black bears and bobcats) made movements that placed them in the proximity of a) mule deer and b) elk more often than expected by chance in northeastern Oregon, 2016-2019. Dots represent point estimates for the selection coefficient for ungulate presence and bars represent 95% confidence intervals. The gray horizontal line at y = 1 indicates indifference toward parturient ungulates; values above this line indicate selection and values below the line indicate avoidance. Cougars and bobcats did not encounter mule deer often enough for models to converge.

**Figure 4:**
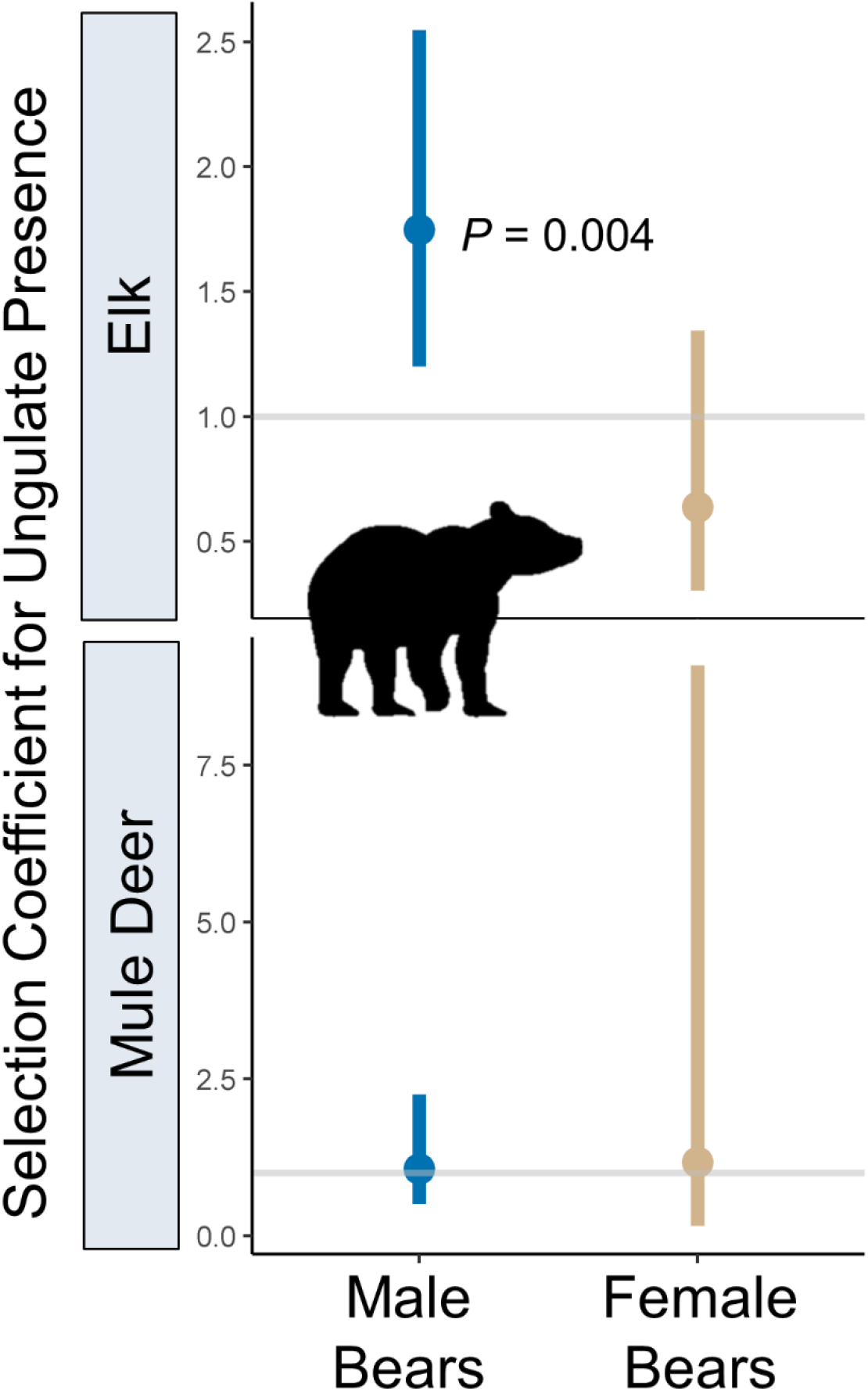
Selection coefficients (exp(β_ungulate presence_)) from step-selection functions assessing whether male and female black bears made movements that placed them in 200-meter proximity of parturient elk more often than expected by chance in northeastern Oregon, 2016-2019. Dots represent point estimates for the selection coefficient for elk presence and bars represent 95% confidence intervals. The gray horizontal line at y = 1 indicates indifference toward parturient elk; values above this line indicate selection and values below the line indicate avoidance.

**Figure 5:**
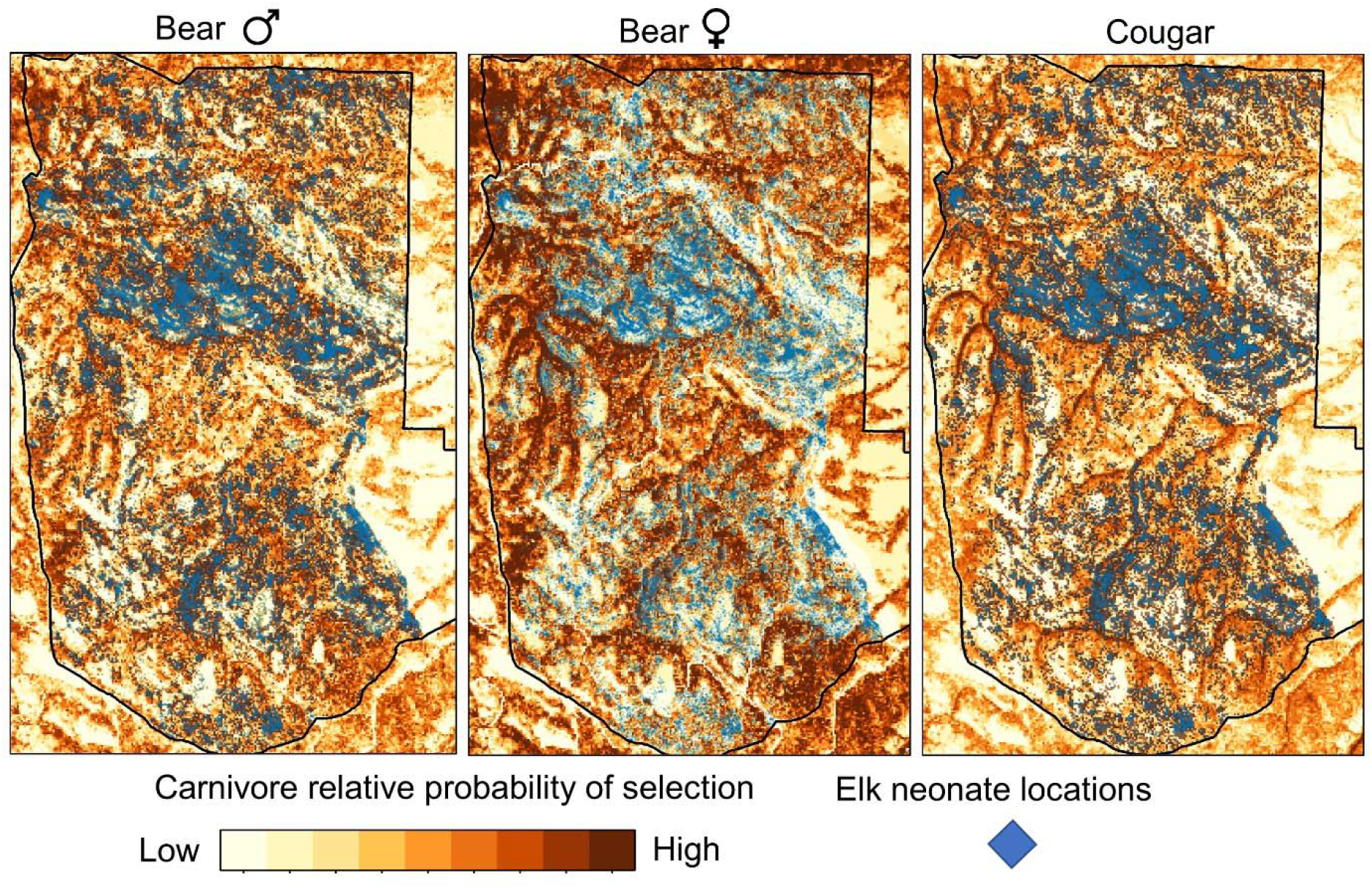
Relative probability of selection for a) male black bears, b) female black bears, and c) cougars in northeastern OR (2016–2019) predicted from step-selection functions (darker shades of brown indicating higher relative probability of selection for each carnivore). In each panel, the GPS locations of telemetered adult female elk in the 30 days post-parturition (blue points) are overlaid on the relative probability of selection maps. The elk locations are displayed identically in each panel but appear darker when they overlap pixels with higher (i.e., darker shades of) relative probabilities of selection for carnivores. The maps are presented as a visual aid to portray how male bears and cougars, but not female bears, select for features of the landscape that overlap with areas used by post-parturient elk. Maps for all species can be found in the Supporting Information (Figures S1–S10).

### Carnivore use of parturition habitat

We hypothesized that the use of predicted parturition habitat by actively-searching predators would track the phenology of the birth pulse such that they would use areas that maximized encounters with neonates when they were most available. Of the species concluded to exhibit active search for neonates (cougars and bears responding to elk calves), linear mixed-models suggested that only male bears followed the expected quadratic pattern for use of parturition habitat coincident with peak parturition date (β_Julian week_ = 0.027, *P* = 0.03; β_Julian week_^2^ = −0.028, *P* = 0.03; Figure 6b) wherein the quadratic terms were supported over a linear term (likelihood ratio test, *P* = 0.03) for the effect of Julian week on use of calving habitat. To assess whether this result could be spuriously driven by bears selecting for parturition habitat for other reasons such as bottom-up forage acquisition, we tested whether female bears with similar vegetation requirements as male bears also responded to parturition habitat but found no support for a quadratic response (likelihood ratio test, *P* = 0.62; Figure 6c). For cougars, use of predicted parturition habitat was also better explained by a linear term for Julian week (likelihood ratio test *P* = 0.32; Figure 6d).

**Figure 6:**
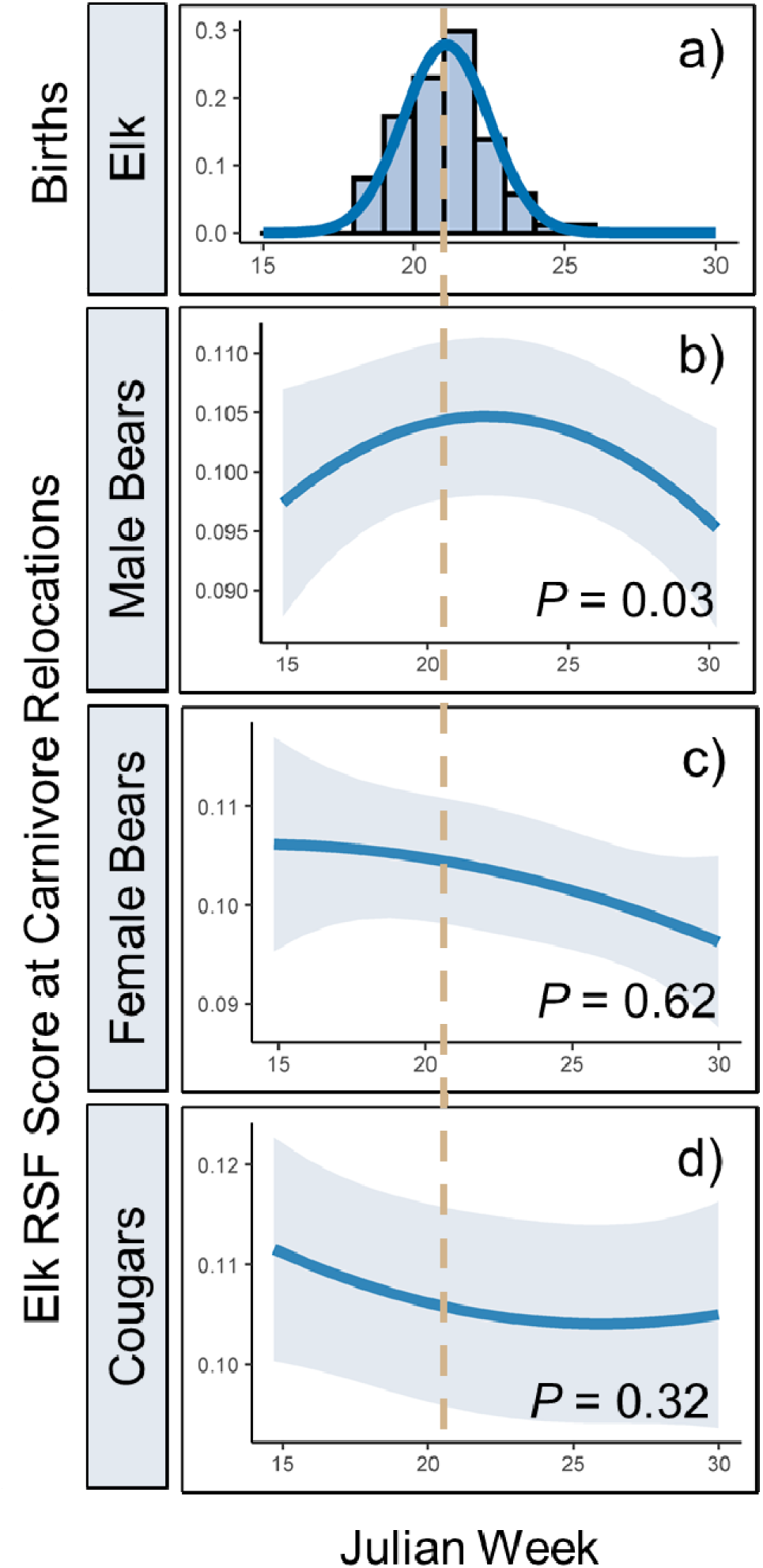
Elk births and carnivore use of predicted elk parturition habitat as a function of Julian week in northeastern Oregon, 2016-2019. a) The density of elk births as a function of Julian week. Bars indicate counts of births per week and the blue curve is a normal distribution fit to the data. The dashed tan line indicates the mean birth date. b-d) Predictions from linear mixed models relating the weekly average resource selection function (RSF) score for parturient elk at carnivore GPS relocations for male bears (b), female bears (c), and cougars (d) as a quadratic function of Julian week. The solid blue lines display the model predictions for carnivore use of elk parturition habitat and the gray shading represents its 95% confidence interval. *P*-values correspond to the outcome of likelihood ratio tests assessing whether the models containing quadratic terms for Julian week were supported over models with only a linear term for Julian week.

## DISCUSSION

We found that of four carnivore species, only two appeared to actively search for elk and none searched for mule deer neonates. The fact that no carnivores encountered parturient mule deer more often than expected by chance was consistent with our predictions that the young of less abundant species would not be targeted (Figure 1b); mule deer are at least 5 times less abundant than elk in this ecosystem (Oregon Department of Fish and Wildlife, unpublished data). This result suggests that mule deer fawn mortalities in this study area (Jackson *et al.* 2021) are likely the result of fortuitous encounters (from the perspective of the predator) and that risk to mule deer from incidental predation may thus depend on the amount of overlap between deer and elk parturition habitat for predators that are primarily searching for elk calves. Our finding that two of the four carnivore species actively searched for elk (the more abundant species) partially aligned with our expectations that a predatory response would be greater toward the more abundant prey species. We also expected the more generalist carnivores (bears and coyotes, which are highly omnivorous) would exhibit a more fluid response to an ephemeral prey source. This hypothesis was supported for bears but not coyotes. The lack of response of bobcats and coyotes to both elk and deer may be explained for several reasons.

First, elk calves are large prey items for mesopredators such as coyotes and bobcats particularly when subject to maternal defense from elk. While mule deer young are much smaller, their rarity on the landscape may not have warranted search behavior. Nonetheless, we can speculate that had mule deer been the more abundant species in this area, we may have observed some level of search behavior by coyotes and/or bobcats. Second, bobcats are efficient predators of small prey while coyotes in this system gain a substantial amount of protein via scavenging (Ruprecht et al. 2021b), both of which potentially reduce the need to pursue neonates. Finally, our sample size of collared bobcats was small (*N* = 6) so results may have been subject to Type II error for that species.

Our analyses to evaluate whether carnivores’ use of elk parturition habitat tracked the phenology of the birth pulse revealed further contrasting behaviors among the predators in our study. Use of elk parturition habitat by male bears was best explained by a model with a quadratic effect of Julian week, with its maximum aligning almost perfectly with the peak of the birth pulse (Figure 6b). This result suggests that male bears exhibited a spatial shift in habitat consistent with an effort to maximize encounters with elk neonates immediately following parturition, which is logical given that bears are limited by a short window in which they can efficiently hunt neonates. This idea is further supported in that the number of encounters between GPS-collared bears and parturient elk in the 30 days post-parturition peaked between 7-10 days after an elk gave birth. By contrast, the quadratic effect was not supported for cougars (Figure 6d). Although previous research has shown that juvenile elk constitute a large fraction of the diets of cougars (Clark *et al.* 2014), several aspects of cougar predatory behavior may explain why cougars did not exhibit a quadratic effect indicative of searching areas used by neonates. First, because elk become solitary to give birth before rejoining the herd several weeks later (Paquet & Brook 2004), it may be inefficient for cougars to target solitary mother-young pairs when they would have access to more individuals by pursuing larger herds of mixed age classes. Such “nursery herds” (Paquet & Brook 2004) would present naïve prey such as yearling elk that may have recently lost maternal guidance, or vulnerable young of the year after the mother-young pair rejoined the herd after parturition. The number of encounters between GPS-collared cougars and parturient elk in the 30 days post birth was greatest around 20 days after an elk gave birth which further supports the idea that cougars pursued calves only after they had matured for several weeks. We were initially concerned that the apparent use of parturition habitat by bears coinciding with the birth pulse could be driven by selection for other dynamic resources such as green vegetation that was correlated with elk parturition habitat, causing a spurious result. However, only male bears exhibited this pattern, and we would expect females to exhibit the same response to bottom-up resources.

Although we are unaware of previous research on search behavior of cougars toward ungulate neonates, our work both aligns and contrasts with patterns described for black bears and coyotes elsewhere. The density of elk neonates in our study area was closer in magnitude to the number of caribou calves in the study by Rayl *et al.* (2018) that determined bears actively searched for neonates than it was to the study by Bastille-Rousseau *et al.* (2011), in which bears opportunistically encountered neonates. Our finding that male bears were much more likely to encounter neonates than were females also aligns with Rayl *et al.* (2015) who found that male bears were more likely to visit caribou calving grounds than were females. Further, several other studies have documented higher predation or meat consumption by male black bears than females (Boertje *et al.* 1988; Jacoby *et al.* 1999), which should be expected given that previous research has shown that larger, male bears require more animal-borne protein to gain weight than do smaller, female bears (Rode, Robbins & Shipley 2001). Our results did not suggest that coyotes actively searched for elk calves which accords with a cause-specific mortality study of elk calves in this region that found coyote predation to be minor (Johnson *et al.* 2019). In other ecosystems, however, coyotes have been implicated as nontrivial sources of mortality for elk calves (Barber-Meyer, Mech & White 2008) although there is mounting evidence that coyote predation on mule deer neonates occurs largely only when small mammal populations are low (Hamlin *et al.* 1984; Hurley *et al.* 2011). We unfortunately did not have sufficient data on occurrence and abundance of alternative prey to assess whether this occurred in our study.

It is important to view the distinction between active search and incidental encounter in light of the effects that predator population dynamics may have on prey. If predators employ active search behavior, then a reduction in predator density may not yield increased neonate survival because neonates spared by that individual become targets for the remaining pool of searching predators (unless prey density is such that predation is limited by handling time or satiation). But if neonate predation occurs because of incidental encounters, then a reduction in predator density benefits a focal prey species by reducing encounter rates both because each prey is less likely to be incidentally encountered per unit time and because predators spend time handling other species. Although population growth rates of ungulates are most sensitive to survival of adult females, this demographic rate is consistently high and stable (Gaillard, Festa-Bianchet & Yoccoz 1998) and many ungulate populations are instead limited by insufficient recruitment due to low neonate survival (Raithel, Kauffman & Pletscher 2007). Thus, knowledge of how different species of carnivores search or encounter different species of prey will be needed to determine the extent to which predator control would be an effective strategy for managing ungulate populations.

A necessary assumption in our study was that predator and adult female ungulate locations within a 200-meter proximity at the time of simultaneous GPS recordings constituted an encounter with an ungulate neonate in the weeks after birth. This assumption was required because we did not have GPS transmitters on neonates and thus assumed the location of the adult female was a reasonable proxy for the location of the neonate. This assumption certainly introduced some amount of data contamination that may have obscured a stronger signal than what we observed. Although our dataset was unique in that it included contemporaneous GPS recordings on four species of carnivores and two species of ungulates, with fix intervals up to three hours on predator GPS collars, we are certain to have missed additional encounters. Another limitation in our study was that we were required to use inference methods to predict some of the parturition events for deer and all parturition events for elk which are subject to error in the exact timing of births. Further, some number of neonates likely died in the days following birth which our analysis could not consider. These factors should only have acted to dilute any potential signals in the data and not cause spurious correlations. Nonetheless, we caution that our results are conservative and should be interpreted with the possibility that Type II error may be present. These issues will be minimized as GPS transmitters become sufficiently lightweight to be placed on neonates and as battery life improves such that more frequent GPS readings can be obtained on both predators and prey.

An emerging frontier in animal ecology is understanding if and how behavior influences population dynamics. Consequently, elucidating how predatory tactics (e.g. active search vs. incidental encounters) affect prey populations should feature prominently in future research. Our results suggest there was a behavioral response by two of the four carnivores toward elk, but no response by any of the carnivores toward mule deer. This result combined with previous research suggests that the foraging tactics by predators in response to a pulsed resource differ by both predator and prey species, are likely ecosystem-specific, and can change dynamically through time as the availability of vulnerable neonates fluctuates.

## ACKNOWLEDGEMENTS

We thank R. Long and M. Bianco for data assistance. We are grateful to the Oregon Department of Fish and Wildlife, the USDA Forest Service Pacific Northwest Research Station, Oregon State University, and the Wildlife Restoration Act for funding and/or logistical support. We thank research technicians and volunteers who assisted with fieldwork.

